# Differential properties of gp120 from HIV-1 group M subtypes A and B

**DOI:** 10.64898/2026.11.09.765515

**Authors:** Yukako C. Tamazawa, Dante I. Suzuki, Yuichiro J. Suzuki, Nataliia V. Shults

**Author notes:** To whom correspondence should be addressed to: Prof. Yuichiro Justin Suzuki, Department of Pharmacology and Physiology, Georgetown University Medical Center, 3900 Reservoir Road NW, Washington DC 20007. contributed equally to this project (Yukako C. Tamazawa, Yuichiro J. Suzuki).

## Abstract

HIV-1 group M subtypes A and B differ substantially in their gp120 sequence. To determine whether these differences contribute to subtype-specific complications, particularly pulmonary arterial hypertension (PAH), rats were injected with gp120 from subtype A or B. Pulmonary artery (PA) remodeling was assessed after four weeks using hematoxylin and eosin (H&E) staining and the Vessel Morphometry Analysis tool by Autona (autona.org). Subtype A gp120 potently increased PA wall thickness and right ventricular hypertrophy. The CD4-binding region of subtype A lacks a stretch of 16 amino acids (V1 *N*-glycosylation site), and experiments showed stronger binding of subtype B to CD4. Subtype B gp120 contains an RGKV sequence, which resembles the RGKR sequence of the rabies virus glycoprotein G that binds to the nicotinic acetylcholine receptor (nAChR). Subtype A gp120 possesses a KDKK sequence instead. Molecular dynamics simulations indicated that both sites are surface-exposed. Although nAChR is expressed in PA cells and could potentially drive the pronounced effects of subtype A, MD simulations estimated that subtype B gp120 binds more strongly to nAChR. We also noted a single amino acid difference immediately preceding the REKR furin cleavage sequence: glutamate in subtype A versus glutamine in subtype B gp120. Simulations predicted that subtype A gp120 interacts more effectively with furin, and enzyme assays demonstrated that subtype A promotes a positive feedback loop more strongly than subtype B. Thus, HIV-1 gp120 subtype A is more potent in promoting PAH, and the mechanism may involve enhancing furin activity to elevate gp120 levels.

## 1. Introduction

HIV is a retrovirus whose viral fusion protein gp160 binds to human CD4 to facilitate fusion, introducing viral RNA into the host cells. CD4, the primary receptor for HIV-1, is largely expressed on the surface of immune cells, and also on vascular endothelial cells (ECs) and smooth muscle cells (SMCs) [Mach, 1997; Schecter, 2001]. gp160 also binds to the co-receptors CCR5 and CXCR4 [Alkhatib, 2009]. gp160 is cleaved into gp120, which contains the receptor binding sites, and gp41, which anchors the complex to the viral membrane, producing free circulating gp120 [Hallenberger, 1992; Richard, 2015]. Circulating gp120 levels of 1 to 8 nM were detected in HIV-infected patients [Cummins, 2010; Oh, 1992]. High gp120 levels occur even in patients receiving antiretroviral therapy (ART) with an undetectable viral load. This protein activates cell signaling events in a variety of cell types, including ECs and SMCs [Del Cornò, 2014; Green, 2014; Hioe, 2011; Schecter, 2001; Yang, 2009]. HIV has infected 92 million people worldwide, and 41 million people are estimated to be currently living with this virus [Deeks, 2015; WHO, 2026].

The HIV classification consists of HIV type 1 (HIV-1) and HIV type 2, and HIV-1 is further stratified into groups M, N, O, and P. Among these, the majority of people living with HIV are infected with HIV-1 group M. Group M is subdivided further into nine subtypes (A-D, F-H, J, and K) [Bbosa, 2019]. The major subtype that is circulating in Western countries, including the U.S., is HIV-1 group M subtype B. Although subtype B accounts for only 10% of HIV infections worldwide [Siemieniuk, 2013; Taylor, 2009], HIV research, which is largely performed in Western countries, is based on this subtype. It is often assumed that the pathobiology of subtype B applies to all other subtypes. By contrast, HIV-1 group M subtype A predominates in Eastern Europe and Central Asia [Nikolopoulos, 2016; Saad, 2006]. Many of the countries affected by HIV-1 subtype A are low- to middle-income countries, and research on this viral subtype has not been rigorous. A shift in the diversity of HIV strains has recently been noted in North America, including the increased presence of subtype A [Siemieniuk, 2013]. Our laboratory previously reported that gp120 from HIV-1 subtypes A and B (which share only 70% identity) elicits different gene expression changes in human pulmonary artery ECs and SMCs [Suh, 2023]. This provides a basis for rigorous investigation into the molecular differences between the actions of HIV components across various HIV-1 subtypes, with the goal of offering tailored therapeutic strategies to prevent and treat HIV-associated complications in patients infected with different subtypes.

Antiretroviral therapies have substantially prolonged the life expectancy of people living with HIV [Davis, 2021; Wandeler, 2016]. As a result, an increased prevalence of non-communicable diseases is now seen in these individuals [Davis, 2021; Ghafouri, 2006; Patel, 2018]. HIV-associated pulmonary arterial hypertension (PAH) occurs in approximately one in 200 HIV-infected patients [Mehta, 2000; Sitbon, 2008; Isasti, 2013; Pellicelli, 2001]. HIV-infected patients have a 2,500-fold increased risk of developing PAH compared to the general population [Bigna, 2015]. A study of 7,648 HIV patients from 2004 to 2005, conducted after the introduction of antiretroviral therapies, showed a prevalence of 0.5% [Sitbon, 2008]. These findings are similar to those from studies conducted before effective antiretroviral therapies were employed, suggesting that these therapies did not alter the prevalence of HIV-PAH and indicating a direct role of HIV components in promoting PAH.

The present study tested the hypothesis that the *in vivo* administration of gp120 from subtypes A and B elicits differential effects on the pulmonary vasculatures in rats. We further utilized molecular dynamics simulations to analyze differential actions of the two gp120 proteins based on their amino acid sequences and three-dimensional structures.

## 2. Materials and Methods

### 2.1. Animal Experiments

Male Fischer CDF rats (Charles River Laboratories, Wilmington, MA, USA) were injected intraperitoneally with 0.4 mg/kg body weight of recombinant gp120 from HIV-1 group M subtype A (isolate 92RW020; Accession Number AAT67478.1; Cat# 40403-V08H) or subtype B (isolate BAL; Accession Number AAA44191.1; Cat# 40404-V08H) from Sino Biological (Paoli, PA, USA) [Suh, 2023]. After 4 weeks, lungs were collected for histology analysis. To assess right ventricular hypertrophy, the right ventricle (RV), left ventricle (LV), and septum (S) were weighed, and the Fulton index was calculated using the equation RV/(LV + S). The Georgetown University Animal Care and Use Committee approved all animal experiments. Our investigation was conducted in accordance with the National Institutes of Health Guide for the Care and Use of Laboratory Animals.

### 2.2. Histology

Lung tissues were immersed in buffered 10% formalin at room temperature, and were then embedded in paraffin. Paraffin-embedded tissues were cut (5 µm thick) and mounted on glass slides, and tissue sections were subjected to hematoxylin and eosin (H&E) staining. Slides were digitally scanned to SVS files.

### 2.3. Vessel Morphometry Analysis

To measure pulmonary artery (PA) wall thickness, the Vessel Morphometry Analysis tool by Autona (autona.org) was used. For quantifications, 20 vessels (30 – 100 µm diameter) were analyzed per animal (the analysis tool identified 200 candidate vessels, from which the user selected 20 for analysis. Values for inner and outer diameters were automatically obtained for each vessel using the analysis system, and the means were calculated. Percent wall thickness values were calculated using the equation (External Diameter – Internal Diameter) / External Diameter by the automated Autona system, using SVS files of H&E images viewed in Aperio ImageScope.

### 2.4. CD4 Binding Assay

To compare the CD4-binding capacities of gp120 from subunit A and subunit B, competition experiments were performed using the RayBio Ligand Binding Assay CD4/IL16 Inhibitor Screening ELISA Kit (Cat# BAH-CD4/IL16; RayBiotech Life, Inc., Peachtree Corners, GA, USA). The assay kit measures the binding of interleukin-16 (IL-16) to CD4 immobilized on a microplate. Bound IL-16 is detected using an anti-IL-16 antibody and a horseradish peroxidase (HRP)-conjugated anti-IgG secondary antibody. HRP activity is quantified with a 3,3’,5,5’-tetramethylbenzidine (TMB) substrate by measuring absorbance at 450 nm using an EMax Plus Microplate Reader (Molecular Devices, LLC., San Jose, CA, USA). The binding of CD4 to the added viral fusion proteins was estimated by competition with CD4/IL16 binding.

### 2.5. Furin Activity Assay

To compare the effects of gp120 from subtypes A and B on furin, the Fluorimetric SensoLyte Rh110 Furin Activity Assay Kit (Cat#: AS-72256; AnaSpec Inc., Fremont, CA, USA), which monitors furin protease activity, was used. The cleavage of the fluorogenic substrate by furin generated the rhodamine 110 fluorophore, which was measured using a POLARstar OPTIMA Fluorescence Microplate Reader (BMG Labtech, Cary, NC, USA). Results at the 120 min time point were reported in relative fluorescence units (RFU).

### 2.6. Molecular Dynamics Simulations

#### 2.6.1. Sequence Data Acquisition, Sequence Alignment, and Variable Region Annotation

HIV-1 envelope glycoprotein gp160 sequences (group M, subtypes A and B) were obtained from the National Center for Biotechnology Information (NCBI) GenBank database: subtype A (accession no. AAT67478.1, 846 aa) and subtype B (accession no. AAA44191.1, 855 aa), based on our previous publication [Suh, 2023]. The gp120 region was defined as residues 1–494 of the Subtype A sequence and residues 1–509 of the subtype B.

The HIV-1 gp160 sequences obtained in this study were aligned against the HXB2 reference strain (GenBank accession no. K03455; Env protein AAB50262.1) using the T-Coffee algorithm within Jalview (version 2.11.5.1; jalview.org) [Waterhouse, 2009], with default parameters. The HXB2 sequence was included in the alignment dataset and used as the positional reference. Alignments were exported in FASTA format and manually inspected to confirm positional homology, particularly across hypervariable regions. Variable loop boundaries (V1–V5) were defined according to HXB2 reference coordinates obtained from the Los Alamos HIV Database (<u>hiv.lanl.gov</u>) [Foley, 2018]. Alignments were visualized using ESPript 3.2 (<u>espript.ibcp.fr</u>) [Robert, 2014] to compare gp160 sequences between Subtypes A and B. Potential *N*-glycosylation sites (sequons) (N-X-S/T motif, where X ≠ P) were identified by manual inspection of the aligned sequences.

#### 2.6.2. Molecular Modeling of gp120 Structures

Three-dimensional structures of gp120 for Subtypes A and B (residues 1–494 and 1–509, respectively) were generated using Boltz-1, an AlphaFold3-like biomolecular structure prediction model, on the Neurosnap platform (<u>neurosnap.ai</u>) [Wohlwend, 2024], with Boltz steering potentials enabled at inference time to reduce steric clashes and enforce stereochemically correct geometry. Model quality was assessed using per-residue predicted Local Distance Difference Test (pLDDT) scores (scale 0–100). The mean pLDDT was 78.9 for Subtype A and 70.4 for Subtype B, indicating moderate-to-high overall confidence, with lower confidence for Subtype B. The models do not include glycans. Final structures were exported as PDB files and visualized using PyMOL (version 2.4.1, Schrödinger, LLC).

#### 2.6.3. RGKV peptide Docking and Molecular Dynamics Simulation

Motifs from α-bungarotoxin (RGKV; PDB: 1KFH), the SARS-CoV-2 spike protein (RRAR; PDB: 6ZGH), rabies virus glycoprotein G (RGKR; PDB: 7U9G), and HIV-1 gp160 Subtypes B (RGKV) and A (KDKK) were extracted with ±15 flanking amino acids using PyMOL; all motifs were surface-exposed in their protein contexts. Each peptide was docked to the α7 nAChR (PDB: 7EKP) using HADDOCK (version 2.4) (wenmr.science.uu.nl/haddock2.4) [Dominguez, 2003; Honorato, 2024] without interface restraints (blind docking), with HADDOCK’s default clustering protocol (FCC-based, default parameters). The highest-scoring pose for each complex was selected for MD simulations (HADDOCK scores ranged from −61.2 to −92.1). Three independent simulations with different initial velocities were run from this pose for each complex using GROMACS version 2019.6 [Abraham, 2015], with the AMBER99SB-ILDN force field and TIP3P water at 300 K. Simulations were run for 5 nanoseconds (ns). Molecular Mechanics/Generalized Born Surface Area (MM-GBSA) binding free energies were calculated using MMPBSA.py [Miller, 2012] from each trajectory.

#### 2.6.4. Docking and Molecular Dynamics Simulations of gp120-Derived Motifs with Furin

Independent 80-ns molecular dynamics simulations were performed for four docking poses per subtype (A1–A4 and B1–B4), each from a distinct docking cluster. For each pose, the peptide and the furin residues within 2.0–3.0 Å of it were extracted to build a reduced system, given computational constraints, and processed with PDBFixer to add hydrogens and complete terminal residues. Simulations were run in OpenMM [Eastman, 2017] with the AMBER14 force field and TIP3P-FB water, using cubic boxes with 1.0-nm padding, neutralized with counterions and 0.15 M NaCl, particle mesh Ewald (PME) electrostatics (1.0-nm cutoff), and constraints on bonds involving hydrogen. Each system was energy-minimized (5,000 iterations), equilibrated for 50 picoseconds (ps) in the constant number of particles, volume, and temperature (NVT) ensemble and 50 ps in the constant number of particles, pressure, and temperature (NPT) ensemble, and then simulated in the NPT ensemble at 310 K and 1 atm using a Langevin middle integrator (friction coefficient, 1 ps⁻¹; time step, 2 femtoseconds) and a Monte Carlo barostat, for a total of 80 ns including equilibration. No positional restraints were applied. Frames were saved every 20 picoseconds. Because the peptide is small relative to furin, RMSD was calculated separately for furin and the peptide after independent least-squares fitting using cpptraj (AmberTools).

#### 2.6.5. MM-GBSA Binding Free Energy Calculations

For each pose with a completed 80-ns trajectory, the OpenMM trajectory was converted to AMBER format using AmberTools (tleap). Binding free energies were calculated with MMPBSA.py [Miller, 2012] using the generalized Born model (igb = 8, 0.15 M salt, default solute and solvent dielectric constants of 1.0 and 80.0). Entropic contributions were not calculated, so ΔG_bind values are enthalpic estimates for relative comparison. For the six poses with completed analysis (A1–A3 and B2–B4), ΔG_bind and its components (van der Waals, electrostatic, polar and nonpolar solvation) were averaged over the analyzed window and compared between subtype A and B by Welch’s t-test. Because the poses represent independent trajectories from a single representative sequence rather than biological replicates, *p* values were interpreted descriptively.

#### 2.6.6. ClusPro Docking of the REKR-Containing Peptides

The furin catalytic domain was obtained from PDB entry 6HZD [Van Lam van, 2019]. The REKR-containing *C*-terminal regions of gp120 were modeled as 10-residue peptides using Boltz-1 (peptide A with REKR motif: KRRVVEREKR, residues 485–494; peptide B with REKR motif: KRRVVQREKR, residues 499–509). The peptides were docked to furin using ClusPro 2.0 (<u>cluspro.bu.edu</u>) [Kozakov, 2017], with furin residues 254–256, 306, 308, and 311–315 (chain A), which line the substrate-binding pocket, set as attraction residues. Docking generated six clusters per subtype (14–632 members for Subtype A; 38–332 members for Subtype B), each characterized by a coefficient weight, docking score, and RMSD relative to the reference cyclic inhibitor c[glutaryl-Arg-Arg-Arg-Lys]-Arg-4-Amba in 6HZD. Because coefficient weights are cluster-level composite scores rather than independent replicate measurements, comparisons between subtypes were interpreted descriptively. The top four poses per subtype, each from a distinct docking cluster, were selected as starting structures for the 80-ns molecular dynamics simulations. These peptides model the cleavage-site motif and its flanking sequence, not full-length gp120.

### 2.7. Statistical Analysis

Means and standard errors of mean (SEM) were computed. Statistical analyses were performed using one-way analysis of variance (ANOVA) with Tukey’s HSD post hoc test. *p* < 0.05 was defined to be statistically significant. Simulation results were compared by Welch’s t-test.

## 3. Results

### 3.1. Effects of gp120 on Pulmonary Vascular Remodeling

Fischer 344 (CDF) rats were injected with recombinant gp120 proteins from HIV-1 group M subtype A or B, and pulmonary vessels were analyzed in H&E-stained lung images. Four weeks after the injection, both gp120 protein had increased pulmonary arterial wall thickness; subtype A gp120 was significantly more effective than subtype B (see Fig. 1A for representative images and Fig. 1B for quantification). Consistently, gp120 from subtype A caused right ventricular hypertrophy (as assessed by the Fulton index), indicating increased pulmonary arterial pressure (Fig. 1C). These results demonstrated that different gp120 structures between subtypes A and B [Suh, 2023] affect the cardiovascular system differently *in vivo*, with subtype A exhibiting more potent effects.

**Figure 1.**
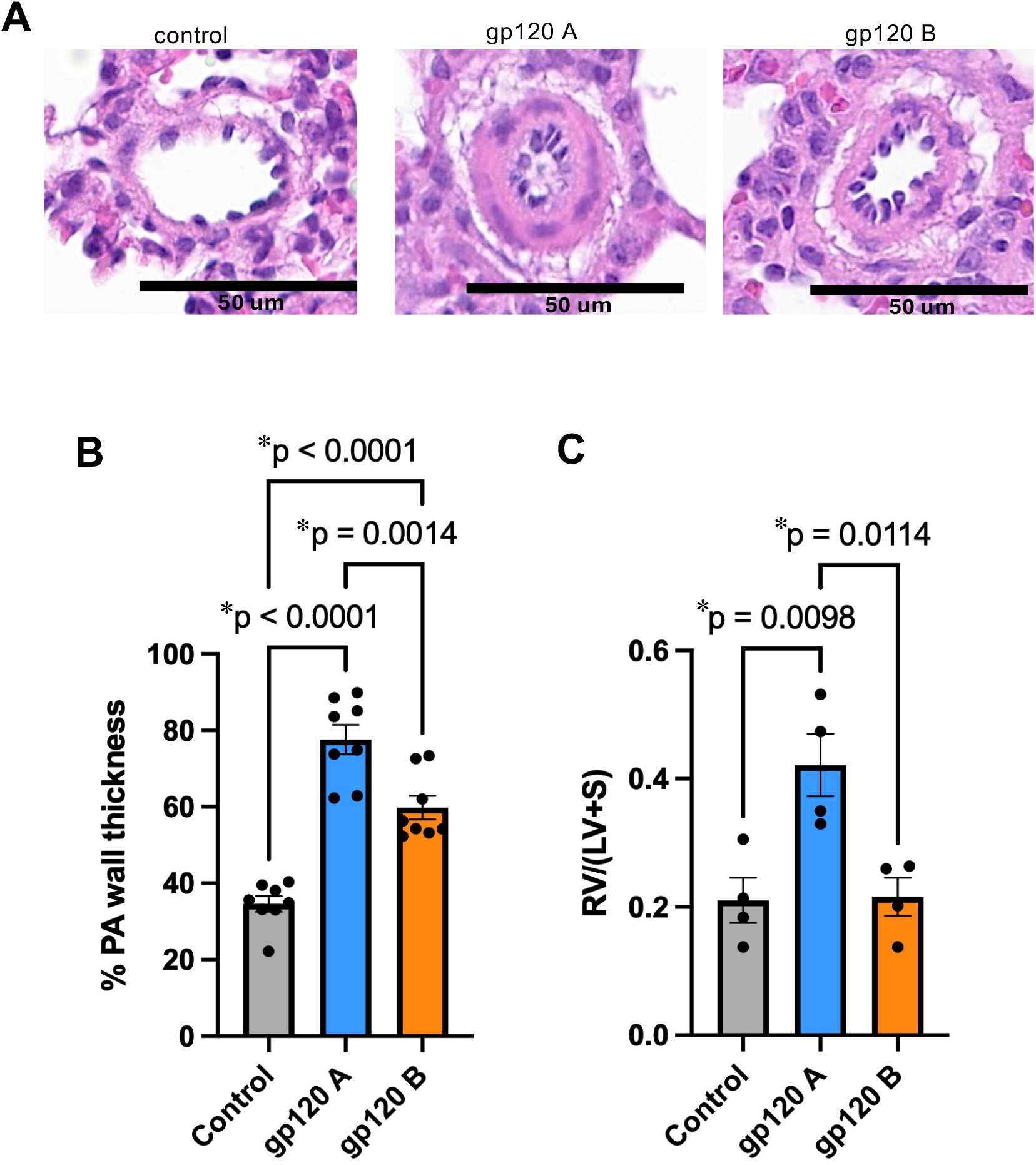
gp120 from subtype A potently promotes the development of pulmonary artery (PA) remodeling and right ventricle (RV) hypertrophy. Fischer 344 (CDF) rats were intraperitoneally injected with gp120 from subtype A (gp120 A) or subtype B (gp120 B). (A) After 4 weeks, lung tissues were immersed in buffered 10% formalin and embedded in paraffin for H&E staining. Representative images are shown. (B) The bar graph represents means ± SEM of % PA wall thickness (n = 8). (C) After 4 weeks of gp120 administration, RV, left ventricle (LV), and septum (S) were weighed and Fulton index, RV/(LV+S), was calculated. The bar graph represents means ± SEM (n = 4). * denotes that the values are significantly different from each other.

### 3.2. Analysis of CD4-binding V1 Loop

To understand how gp120 from subtype A may exert different effects from subtype B, the aligned primary sequences of gp120 proteins from our prior publication [Suh, 2023] were carefully analyzed.

The most notable difference between gp120 sequences of the two subtypes is that the CD4-binding V1 region of subtype A lacks a stretch of 16 amino acids (Fig. 2A). The V1 loop is flexible and a major target of *N*-linked glycosylation. Fig. 2B shows the modeled three-dimensional structures of both gp120 subtypes, depicting the structural differences in this critically exposed region. In these differential V1 loop structures, subtype B contains an additional *N*-linked glycosylation motif (NTT sequence) that is absent in subtype A (Fig. 2C). These sequence and structural differences in the V1 region raise the possibility that subtype-specific gp120 features influence CD4 engagement [Fox, 1997; Kolchinsky, 2001; Wang, 2016], because the V1 loop lies close to the CD4-binding site and undergoes conformational rearrangement upon receptor binding [Guttman, 2012; Kwong, 1998]. We tested the hypothesis that gp120 from subunit A, which possesses fewer glycosylation sites within the V1 region, possesses less ability to bind to CD4. To test this, we used an IL-16/CD4 competition assay, in which lower absorbance values indicate greater displacement of IL-16 from CD4 and therefore stronger CD4 binding by the competitor. At 100 nM, gp120 from both subtypes reduced IL-16/CD4 binding, with subtype B being more effective (Fig. 3A). Dose–response experiments (0–200 nM) also showed that subtype B has more potent effects than subtype A in reducing IL-16/CD4 binding, whereas the SARS-CoV-2 spike protein S1 subunit, used as a negative control, had no effect (Fig. 3B). These results indicate that subtype B gp120 binds CD4 more strongly than subtype A, which is consistent with our molecular analysis but does not seem to account for the superior ability of subtype A to promote vascular remodeling.

**Figure 2.**
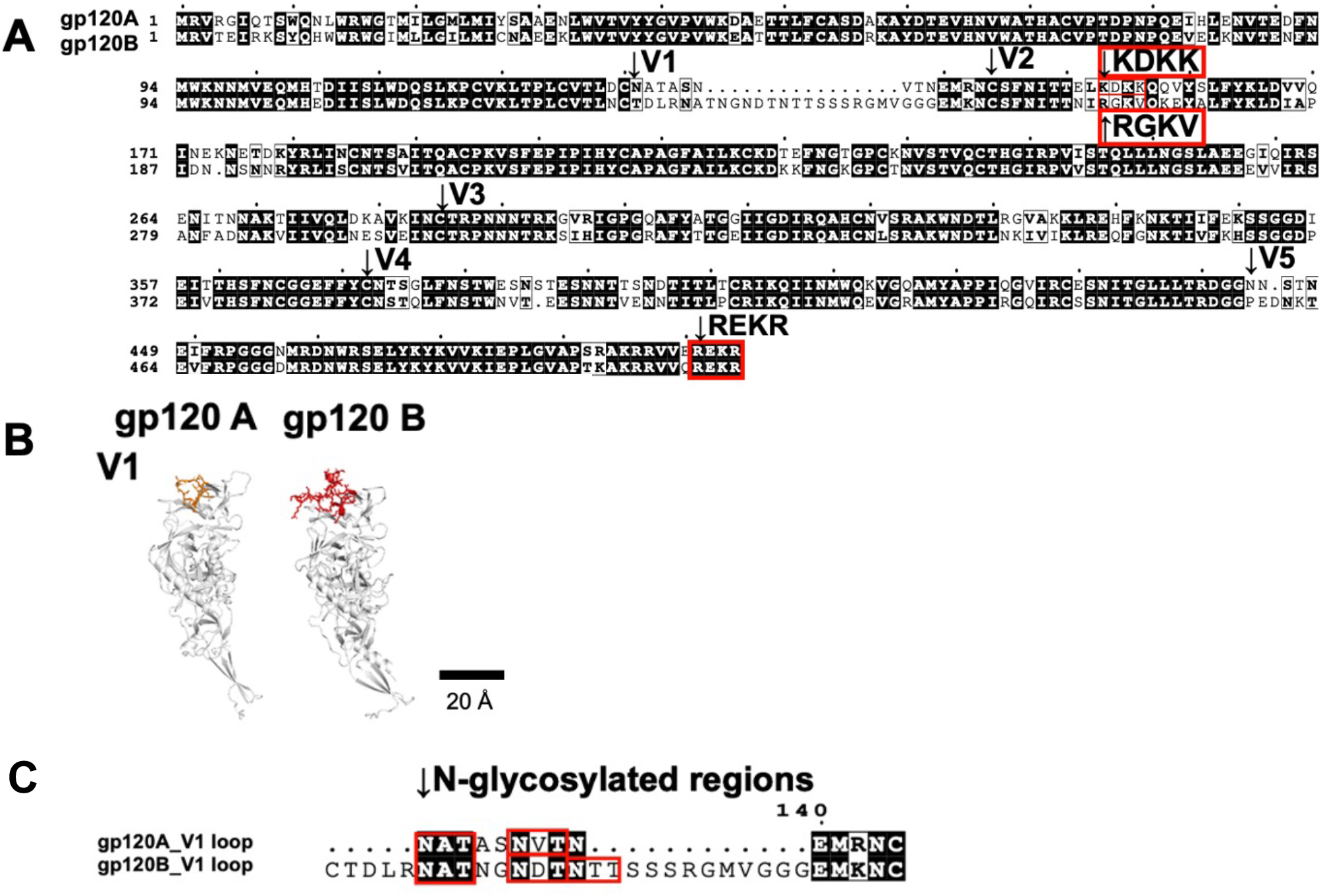
Sequence alignment and structural modeling of gp120 from HIV-1 group M subtypes A and. **B.** (A) Full-length gp120 alignment of HIV-1 group M Subtypes A and B, highlighting the variable loops (V1–V5). The red box indicates the nAChR-binding-like motif (RGKV/KDKK) and the furin cleavage motif (REKR), which were further characterized in later chapter. (B) 3D-modeling of gp120 Subtype A (left, V-loop; orange) and Subtype B (right, V-loop; red). (C) Detailed alignment of the V1-loop showing *N*-glycosylation motifs (N-X-S/T, where X ≠ P). Subtype B carries an additional NTT site in the V1 loop compared with Subtype A.

**Figure 3.**
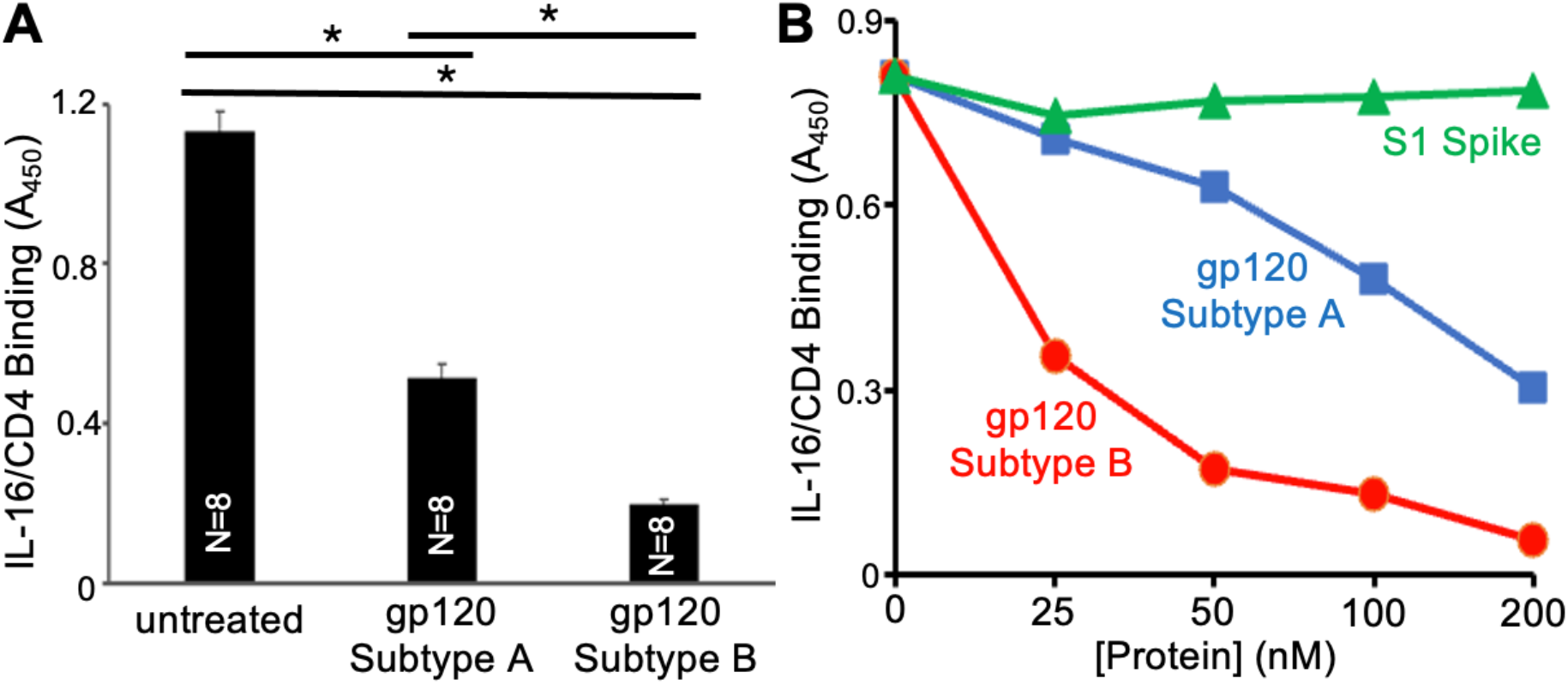
CD4 binding affinity of gp120 from subtypes A and B measured by IL-16 displacement assay. Gp120 from subtype A or B were added to Subtype B gp120 demonstrates significantly higher CD4 binding affinity compared with Subtype A. Lower absorbance indicates greater displacement of IL-16 and thus higher CD4 binding affinity. (A) gp120 from subtypes A and B (100 nM) were incubated with IL-16 in CD4-coated microplate wells. The interference of IL-16 binding to CD4 by gp120 was monitored using IL-16 antibody and HRP-linked secondary antibody while measuring the product of TMB at A_450_. The bar graph represents means ± SEM (n = 8). * denotes that the values are significantly different from each other at *p* < 0.05. (B) Dose-response curves showing competition for IL-16/CD4 binding by gp120 Subtype A (blue squares), gp120 Subtype B (red circles), and SARS-CoV-2 spike protein S1 as a negative control (green triangles).

### 3.3. Analysis of Nicotinic Acetylcholine Receptor (nAChR)-Binding Motif

It has been reported that gp120 from subtype B contains an RGKV motif in the V2 region [Capó-Vélez, 2018; Ballester, 2012] that matches a sequence in α-bungarotoxin [Samson, 2002] and resembles the corresponding motif in rabies virus glycoprotein G (RGKR) [O’Brien, 2024], both of which bind to nAChR. The SARS-CoV-2 spike protein also carries a related basic motif (RRAR) [Changeux, 2020; Hoffmann, 2020; Oliveira, 2021]. However, gp120 from subtype A lacks this RGKV motif, and the corresponding region of subtype A contains KDKK instead. These nAChR-binding sequences of the five proteins are shown in Fig. 4A. We created three-dimensional models of α-bungarotoxin (Fig. 4B), the SARS-CoV-2 spike protein (Fig. 4C), the rabies virus glycoprotein G (Fig. 4D), gp120 from HIV-1 subtype B (Fig. 4E), and subtype A (Fig. 4F). The modeling results revealed that nAChR-binding motifs of all five proteins are surface-exposed.

**Figure 4.**
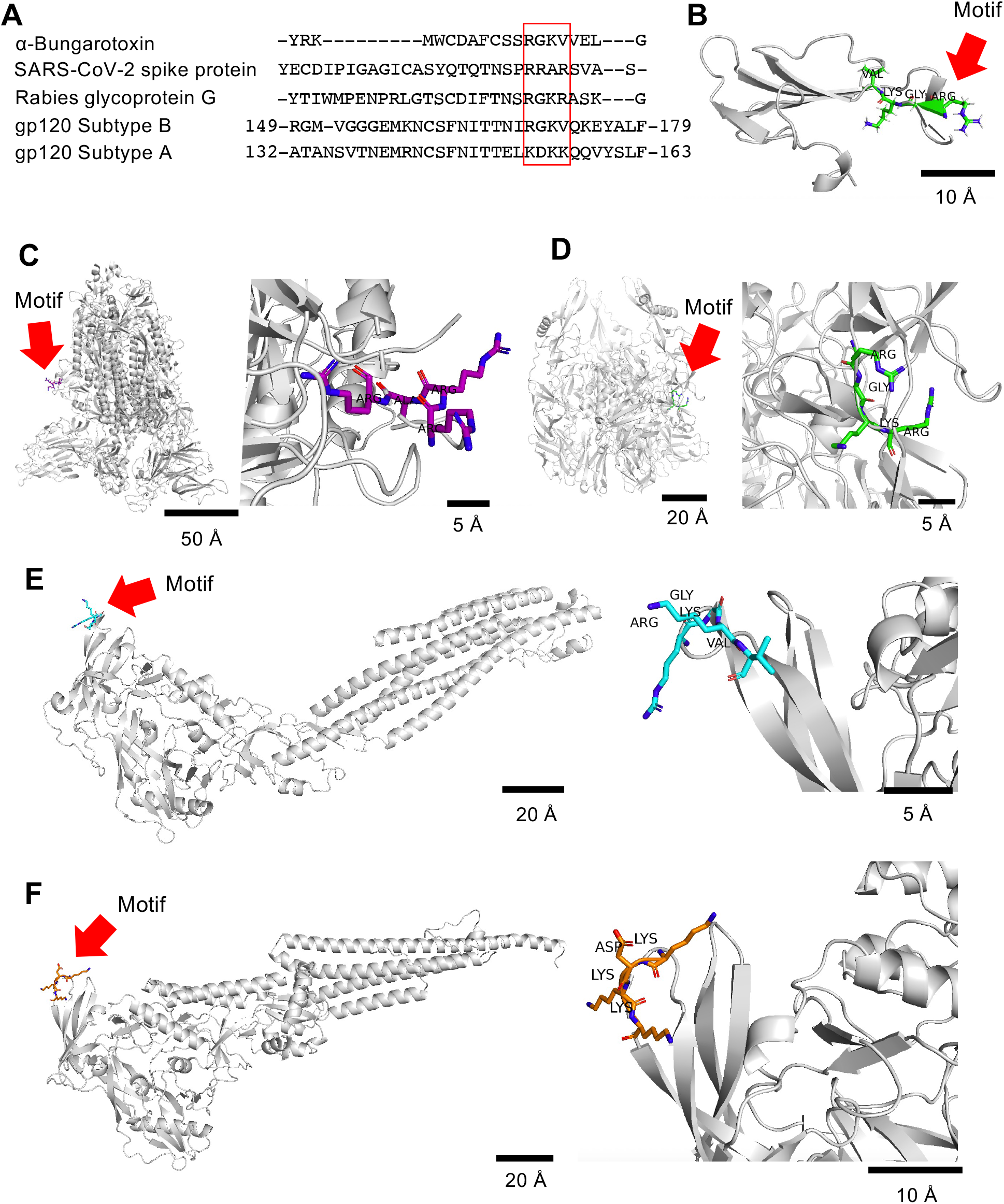
nAChR-binding motifs. (A) Sequences containing nAChR-binding motifs of the five proteins, indicated in the red box. (B-F) Three-dimensional models with nAChR-binding motifs indicated for: (B) α-Bungarotoxin (with RGKV), (C) SARS-CoV-2 spike protein (with RRAR), (D) rabies virus glycoprotein G (with RGKR), (E) HIV-1 gp120 subtype B (with RGKV), and (F) HIV-1 gp120 subtype A (with KDKK).

To compare how the gp120 motifs from the two subunits interact with the nAChR, we performed 5-ns MD simulations of the α7 nAChR with gp120 peptides from subtype A (KDKK; residues 138–171) and subtype B (RGKV; residues 154–187) (sequences shown in Fig. 5A). Results of these simulations depicting the interactions between nAChR with α-bungarotoxin (Fig. 5B), SARS-CoV-2 spike (Fig. 5C), rabies glycoprotein G peptides (Fig. 5D), gp120 from subtype B (Fig. 5E), and gp120 from subtype A (Fig. 5F) are shown. MM-GBSA binding free energies, calculated from the 2–5 ns portion of each trajectory, were less favorable for subtype A than for subtype B (Table 1). These results suggest that, in this simulation framework, the subtype B peptide is predicted to bind nAChR more favorably than the subtype A peptide.

**Fig. 5.**
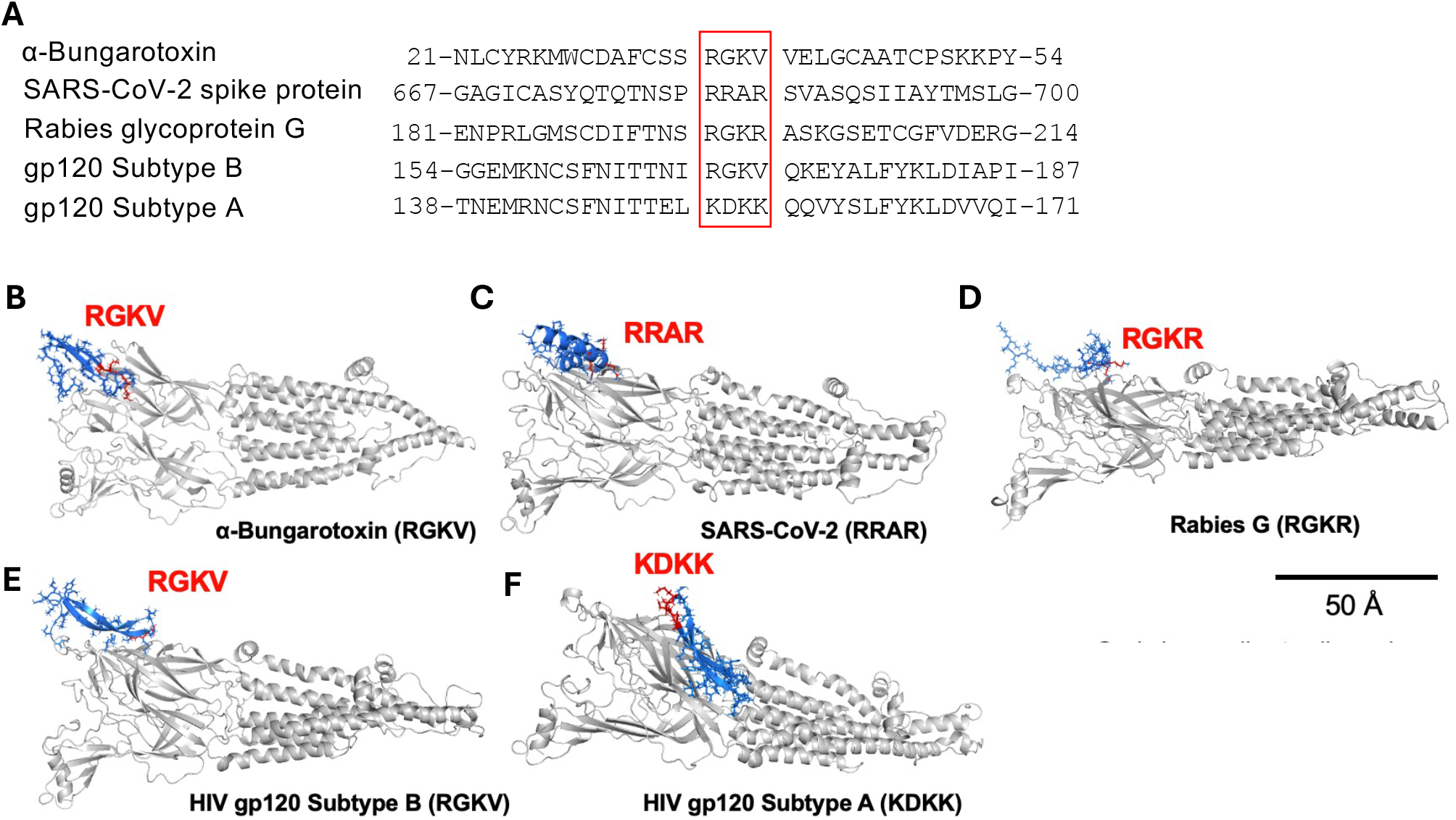
Molecular dynamics simulations of nAChR–peptide complexes. (A) Peptide sequences used in the simulations. nAChR-binding motifs are shown in the red box. (B) Sequences Representative structures from 5-ns molecular dynamics simulations of α7 nAChR with (B) α-bungarotoxin, (C) SARS-CoV-2 spike peptide, (D) rabies virus glycoprotein G, (E) HIV-1 gp120 subtype B, and (F) HIV-1 gp120 subtype A (KDKK). The α7 nAChR structure is indicated in gray, peptide backbones are in blue, and motif residues are in red. Scale bar, 50 Å.

**TABLE 1.**
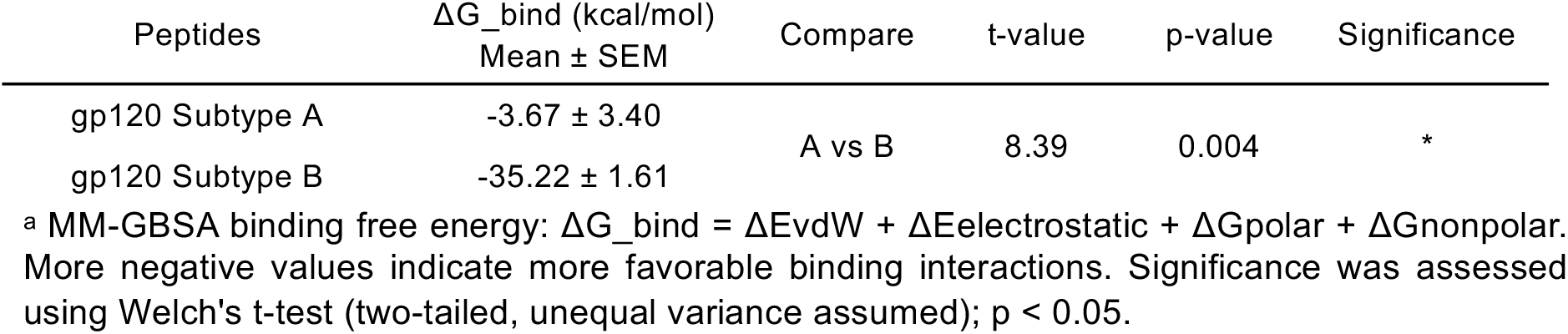
5-ns MD Simulation and MM-GBSA Analysis of Peptide-nAChR Interactions (n = 3)

| Peptides | ΔG_bind (kcal/mol)<br>Mean ± SEM | Compare | t-value | p-value | Significance |
|---|---|---|---|---|---|
| gp120 Subtype A | -3.67 ± 3.40 | A vs B | 8.39 | 0.004 | * |
| gp120 Subtype B | -35.22 ± 1.61 |  |  |  |  |

<sup>a</sup> MM-GBSA binding free energy: ΔG\_bind = ΔEvdW + ΔEelectrostatic + ΔGpolar + ΔGnonpolar. More negative values indicate more favorable binding interactions. Significance was assessed using Welch's t-test (two-tailed, unequal variance assumed); p < 0.05.

### 3.4. Analysis of Furin Motif

By analyzing the primary sequence of gp120 from subtypes A and B [Suh, 2023], we noted that, while gp120 proteins of both subtypes contain the conserved REKR furin cleavage motif (P4–P1), the residue immediately preceding the REKR sequence (P5) differs: glutamic acid (E) in subtype A and glutamine (Q) in subtype B. We hypothesized that the REKR sequence retained at the *C*-terminus of gp120 after furin-mediated cleavage might continue to be recognized by furin as a product-like pseudo-substrate in a subtype-specific fashion.

The 16-residue peptides spanning the cleavage site were used for docking and simulation: peptide A (KRRVVE**REKR**) and peptide B (KRRVVQ**REKR**; Fig. 6). To compare REKR–furin binding between subtypes, the peptide A and peptide B were independently docked to the furin catalytic domain using ClusPro 2.0, and the resulting poses were compared with the co-crystallized cyclic inhibitor c[glutaryl-Arg-Arg-Arg-Lys]-Arg-4-Amba (PDB: 6HZD). Four independent 80-ns molecular dynamics simulations were subsequently performed per subtype, each starting from one of the top four ClusPro-derived docking poses (Table 2). Four interface metrics were evaluated, and all showed statistically significant differences between subtypes (Welch’s t-test; Table 2). Backbone RMSD was significantly lower for Subtype A than for Subtype B (Table 2), indicating that peptide A maintained a more conformationally stable binding pose within the furin catalytic site throughout the 80-ns trajectory, whereas peptide B exhibited greater structural deviation from the initial docked conformation. H-bond occupancy between the REKR motif and furin was markedly higher for subtype A than for subtype B (Table 2), indicating that the subtype A REKR motif sustained more persistent hydrogen bonding with the furin binding site across the simulation, whereas subtype B exhibited more intermittent hydrogen bonding. Both the peptide and the furin binding site showed significantly larger SASA in the subtype A complex than in the subtype B complex (Table 2). Because larger SASA is not typically associated with tighter binding, this metric is reported descriptively. Taken together, all four interface-metrics differed significantly between subtypes, and RMSD and H-bond occupancy indicated that the REKR motif of subtype A engages the furin catalytic site with greater structural stability and more persistent hydrogen bonding than that of subtype B.

**Figure 6.**
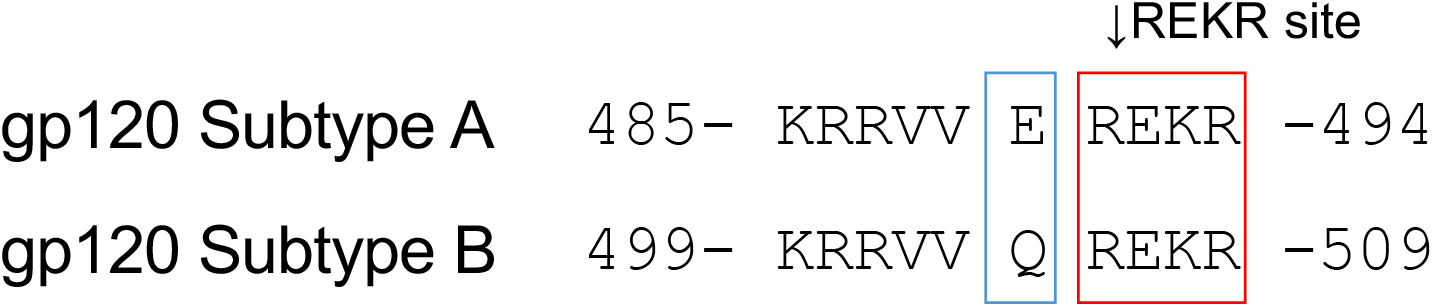
Sequence of the furin recognition site at the gp120 C-terminus. The REKR motif (P4–P1; red box) is conserved, whereas the residue at P5 differs: glutamic acid (E) in Subtype A and glutamine (Q) in Subtype B (blue box). The arrow marks the cleavage site. Numbers indicate residue positions in full-length gp120.

**TABLE 2.** gp120 peptide–furin molecular dynamics simulation analysis (80-ns, n= 4)

| Metric | Subtype A<br>Mean±SEM | Subtype B<br>Mean±SEM | t-value | p-value | Significance |
|---|---|---|---|---|---|
| RMSD (Å) | 1.581 ± 0.367 | 3.915 ± 0.713 | -2.910 | 0.0380 | * |
| H-bond occupancy (%) | 94.08 ± 3.43 | 47.10 ± 7.68 | 5.588 | 0.0045 | * |
| REKR motif SASA (P4–P1) (Å²) | 1824.78 ± 29.18 | 1710.61 ± 19.09 | 3.274 | 0.021 | * |
| Furin binding-site SASA (Å²) | 2146.60 ± 169.07 | 1470.84 ± 96.07 | 3.475 | 0.0193 | * |

Four 80-ns simulations were performed per subtype, each starting from one of the top four ClusPro poses (each from a distinct docking cluster). Data represent means ± SEM; \*p < 0.05 (Welch’s t-test, uncorrected). For pose A3, the analysis window was 68–80 ns (Section 2.6).

The MM-GBSA energy component analysis of REKR–furin interactions revealed that only the van der Waals term differed significantly between subtypes, being more favorable for subtype A (Table 3), which may reflect more favorable shape complementarity within the furin pocket.

**TABLE 3.** 80-ns MD MM-GBSA analysis for gp120 Subtype A and Subtype B (n = 3)

|  | Subtype A<br>Mean±SEM | Subtype B<br>Mean±SEM | A−B | t | df | p |
|---|---|---|---|---|---|---|
| *E*<sub>vdW</sub> (kcal/mol) | −30.84 ± 3.10 | −15.68 ± 3.69 | -15.16 | -3.14 | 3.89 | 0.036 |

Consistent with molecular simulations predicting that subtype A gp120 interacts more effectively with furin than subtype B, furin enzyme assay experiments demonstrated that the subtype A protein promotes a positive feedback loop more strongly than does subtype B. As shown in Fig. 7, a fluorogenic SensoLyte Rh110 Furin Activity Assay revealed that both gp120 subtype A and subtype B significantly increased furin enzymatic activity compared to untreated controls, with subtype A producing significantly greater furin activation than subtype B (Fig. 7A). Dose-response analysis confirmed that this difference was maintained across the tested concentration range (0–300 nM), with subtype A (blue) enhancing furin activity more effectively than subtype B (red) (Fig. 6B). These results identify a molecular mechanism by which gp120 from subtype A exhibits superior effects to those of subtype B.

**Figure 7.**
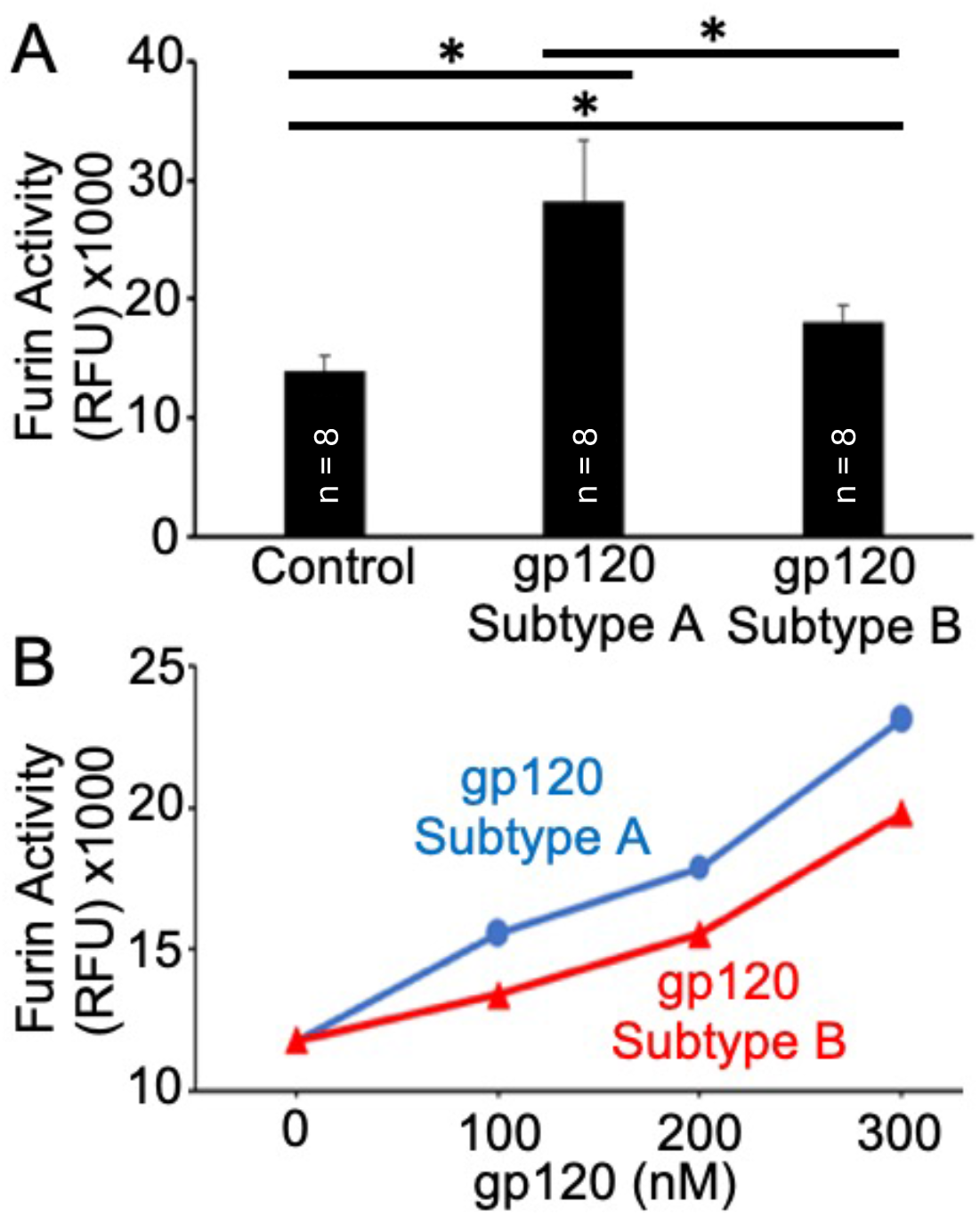
gp120 Subtype A preferentially activates furin enzymatic activity. (A) gp120 Subtype A activated furin activity. Human furin was treated with gp120 Subtype A or B, and its enzymatic activity was monitored using the SensoLyte Rh110 Furin Activity Assay. The bar graph represents the mean ± SEM (n = 8). Statistical significance was determined by one-way ANOVA with Tukey’s post-hoc test (*p < 0.05). (B) Dose-response curve showing dose-dependent activation of furin by gp120 Subtype A (blue) compared with gp120 Subtype B (red). Subtype A exhibits higher furin activation across all tested concentrations (0–300 nM).

## Discussion

Remarkable advances in HIV research have turned this once-deadly disease into a manageable condition [Fauci, 2020]. This, however, has led to an increased incidence of HIV-associated complications such as pulmonary hypertension [Mehta, 2000; Sitbon, 2008; Isasti, 2013; Pellicelli, 2001]. While HIV-1 group M subtype B affect only 10% of people infected with HIV, because this subtype is predominant in leading countries in medical research such as the United States, those in Western Europe, and Japan, much of the information about HIV biology have been generated by studying the subtype B virus [Siemieniuk, 2013; Taylor, 2009]. Many other HIV subtypes, however, are circulating through the world [Bbosa, 2019; Siemieniuk, 2013; Taylor, 2009]; nevertheless, it has been assumed that the information provided for the subtype B virus applies to other types, and the treatment strategies have not been subtype-specific.

In our previous publication, we compared the gp120 sequences of subtypes A and B and found that they share only 70% identity [Suh, 2023]. Treatment of cells with gp120 proteins from subtypes A and B resulted in differential gene expression [Suh, 2023]. The present study demonstrates that these two gp120 proteins can elicit different effects *in vivo*. The administration of gp120 to rats promoted pulmonary vascular remodeling, a hallmark of pulmonary hypertension, with subtype A being most effective. These experimental results suggest that people living with HIV in countries affected by subtype A may have a higher prevalence of HIV-associated PAH, warranting further epidemiological studies.

Building on the differential effects of subtypes A and B of gp120 observed in our previous work [Suh, 2023] and the current study, we employed molecular dynamics simulations and biochemical assays to identify the specific gp120 structural features responsible for these distinct biological activities.

A comparison of the amino acid sequences of gp120 molecules from subtypes A and B clearly highlights the lack of an amino acid stretch in subtype A. These 16 amino acids in the V1 loop bind to the major host cell receptor for HIV, CD4 [Kolchinsky, 2001; Wang, 2016]. The binding of gp120 to CD4 is known to be determined by the extent of glycosylation within the gp120 molecule [Guttman, 2012; Kolchinsky, 2001; Li, 1993], and the absence of these amino acids in the subtype A protein results in fewer glycosylation sites in this region. Thus, it is possible that having fewer glycosylation sites reduces the ability to bind to CD4; indeed, our experiments demonstrated that gp120 from subtype B exhibits a greater capacity to compete with IL-16 for CD4 binding. Thus, gp120 subtype B is more potent in eliciting CD4 binding, and the gp120-CD4 signaling pathway may not be a good candidate for the mechanisms of subtype A-induced pulmonary vascular remodeling.

Interestingly, it has been shown that viruses such as HIV [Capó-Vélez, 2018; Ballester, 2012] and SARS-CoV-2 [Changeux, 2020; Hoffmann, 2020; Oliveira, 2021] contain motifs resembling those found in the rabies virus glycoprotein G, which binds to nAChR and causes neurotoxicity [O’Brien, 2024]. These motifs could be responsible for neurological complications that have been documented in HIV infection [Diaz, 2025] as well as COVID-19 [Pang, 2024]. The reported nAChR-binding motif in gp120 from subtype B is the RGKV sequence, which resides at amino acids positions 154 through 187. While no prior studies have investigated this feature in subtype A or any other HIV subtypes, our analysis reveals that subtype A gp120 lacks this sequence. In these amino acid positions, the subtype A protein contains the KDKK sequence. Thus, gp120-nAChR signaling may not be a good candidate for the mechanism of subtype A-induced pulmonary vascular remodeling.

The next difference we noticed between the gp120 molecules of subtypes A and B was a single amino acid immediately preceding the conserved furin-cleavage sequence. Interestingly, at this position, subtype A has a negatively charged Glu residue, whereas subtype B has an uncharged Gln.

Membrane fusion of HIV with host cells requires proteolytic cleavage of the gp160 protein into gp120 and gp41. The furin protease cleaves at the amino acid sequence Arg-X-Lys/Arg-Arg [Hallenberger, 1992]. In cells, furin accumulates in the Golgi apparatus and can also transported to the plasma membrane [Braun, 2019]. We propose that in viruses associated with severe vascular pathologies, such as HIV and SARS-CoV-2, extracellular furin may process circulating viral membrane proteins that contribute to these complications [Suzuki, 2020]. HIV-infected individuals exhibit 1 - 8 nM of free gp120 in their serum [Cummins, 2010; Oh, 1992]. Observations that furin activity is enhanced by its own product introduce a novel concept: a positive feedback mechanism that may amplify the production of free gp120.

Consistent with molecular simulations predicting that subtype A gp120 interacts more effectively with furin than subtype B, our experimental results revealed that subtype A gp120 strongly enhances furin’s proteolytic activity. This furin-dependent mechanism may increase circulating gp120 levels, potentially accounting for the observed *in vivo* differences between these HIV-1 subtypes and serving as a target for subtype-specific therapies.

